# Gaze and Arrows: does the Gaze Following Patch in the posterior temporal cortex differentiate social and symbolic spatial cues?

**DOI:** 10.1101/2024.01.15.575547

**Authors:** Marius Görner, Hamidreza Ramzanpour, Peter Dicke, Peter Thier

**Affiliations:** Hertie Institute for Clinical Brain Research, Cognitive Neurology Laboratory, Tübingen, Germany; GTC of Neuroscience, Tübingen, Germany; IMPRS for Cognitive and Systems Neuroscience, Tübingen, Germany; Werner Reichardt Centre for Integrative Neuroscience, Tübingen, Germany; Centre for Vision Research, York University, Toronto, Canada

## Abstract

The *Gaze Following Patch* (GFP) is located in the posterior temporal cortex and has been described as a cortical module dedicated to processing other people’s gaze direction in a domain-specific manner. Thus, it appears to be the neural correlate of Baron-Cohen’s Eye-Direction Detector (EDD) which is one of the core modules in his *Mindreading System* - a neurocognitive model for the *Theory of Mind* concept. Inspired by Jerry Fodor’s ideas on the modularity of the mind, Baron-Cohen proposed that, among other things, the individual modules are domain-specific. In the case of the EDD this means that it exclusively processes eye-like stimuli to extract gaze direction and that other stimuli, that may carry directional information as well, are processed elsewhere. If the GFP is indeed EDD’s neural correlate it must meet this expectation. To test this, we compared the GFP’s BOLD activity during gaze-direction following with the activity during arrow-direction following. Contrary to the expectation based on the assumption of domain specificity we did not find a differentiation between gaze and arrow-direction following. In fact, we were not able to reproduce the GFP as presented in previous studies. A possible explanation is that in the present study – unlike previous work –, the gaze stimuli did not contain an obvious change of direction that represented a visual motion. Hence, the critical stimulus component responsible for the identification of the GFP in previous experiments might have been visual motion.

## Introduction

The *Gaze Following Patch* (GFP) is a circumscribed region in the posterior part of the temporal cortex which was discovered in healthy human subjects that participated in fMRI experiments in which the task was to use the gaze direction of a demonstrator to identify a target object among distractors.^1–5^ In contrast with the respective control condition, iris-color mapping in which the observer had to shift gaze to an object whose color corresponded to the color of the demonstrator’s iris, the gaze-following condition yielded a significantly larger BOLD response within the GFP. This preference for gaze direction suggests that the GFP might be the neural realization of Baron-Cohen’
ss Eye Direction Detector (EDD).^6^ Being an integral component of his mind-reading model, Baron-Cohen proposed the EDD to be domain specific^7^. This implies that it exclusively processes eye-like stimuli and forwards information on eye-direction to downstream modules to form a Theory of Mind (ToM), a concept that captures the assignment of desires, beliefs and intentions to another person. Electrophysiological studies in non-human-pimates (NHP) that investigated the response preferences of individual neurons in a presumably homologous brain area in the superior temporal sulcus (STS) seemed to be in line with the assumed domain-specificity of the GFP, in accordance with the central tenet of Baron-Cohen concept.^8^

In his work, Baron-Cohen suggested that in conjunction with shape and contrast patterns, the visual motion signal inevitably yoked with the view of an eye movement plays a crucial role in the detection of eye-gaze stimuli. Hence, under the assumption that the GFP indeed corresponds to Baron-Cohen’
ss EDD its location in a brain region known for its role in visual motion processing appears plausible. However, one may wonder to which extent the GFP is indeed selective for motion of the eyes, a selectivity to be met in order to satisfy the assumption of domain-specificity. In fact a critical examination of this question is still lacking. This is why we embarked on the current fMRI study in which we compared the BOLD activity patterns resulting from contrasting gaze-direction following and iris-color mapping with an analogous contrast: arrow-direction following versus arrow-color mapping. We predicted that the GFP should remain silent in the arrow-contrast condition if it met the premise of domain specificity.

In this study we tested 20 healthy human participants using the same stimuli as in Marquardt et al.^4^ with an important modification: while individual trials in the original version of the study always started with the demonstrator’s overt gaze directed towards the participant followed by a second frame depicting the demonstrator looking towards the target object, in the current study the initial part was replaced by a blank screen. This bears the consequence that no overt gaze shift is seen as the blank screen is directly followed by the demonstrator’s gaze directed towards the target (see Figure 1). Hence, the spatiotemporal discontinuity (apparent motion) of the two views of the eyes in the original version which created the impression of a saccadic gaze shift was absent. This modification was necessary to allow fully analogous sequences in the gaze and arrow conditions, since in two-dimensional views of arrows there is no equivalent orientation to the gaze being directed towards the participant, while still being recognizable as arrows. The presence of apparent motion^9,10^ in the original version of the paradigm had an important consequence; by contrasting the gaze-following condition with the respective control condition - eg. iris-color mapping - we implicitly contrasted a stimulus component that comprised a motion event (the gaze-shift) with a component that comprised a color change requiring to ignore the motion event. Or, to put it differently, in the gaze-following condition, the stimulus component that was behaviorally relevant, i.e. the gaze-direction, was intrinsically linked to visual motion while in the iris-color condition it was not. It has been shown that both electrophysiological and BOLD signatures of visual motion in parts of the posterior STS are boosted whenever motion patterns are behavioral relevant.^11,12^ This raises the question if the human GFP as described by Marquardt et al. and other studies as well as the monkey’s homologue may be the result of this implicit contrast between behaviorally relevant motion and a control condition, lacking behaviorally relevant motion cues. Therefore, in the current study, the first question was whether we are able to reproduce the GFP despite the absence of any motion information provided by the stimulus. Second, we compared the activity patterns emerging from the contrast *gaze-direction* > *iris-color* with those resulting from the contrast *arrow-direction* > *arrow-color* as well as with the location of the GFP as reported by Marquardt et al^4^. We expected that if the GFP is domain-specific and does not depend on visual motion we would not find overlapping activation between the two contrasts at the expected location of the GFP at a given statistical threshold. Moreover, for each condition we estimated the hemodynamic response functions based on the GFP region of interest (ROI) reported by Marquardt et al. and based on a ROI stemming from a search for *visual motion* in the Neurosynth^13^ database.

**Figure 1.**
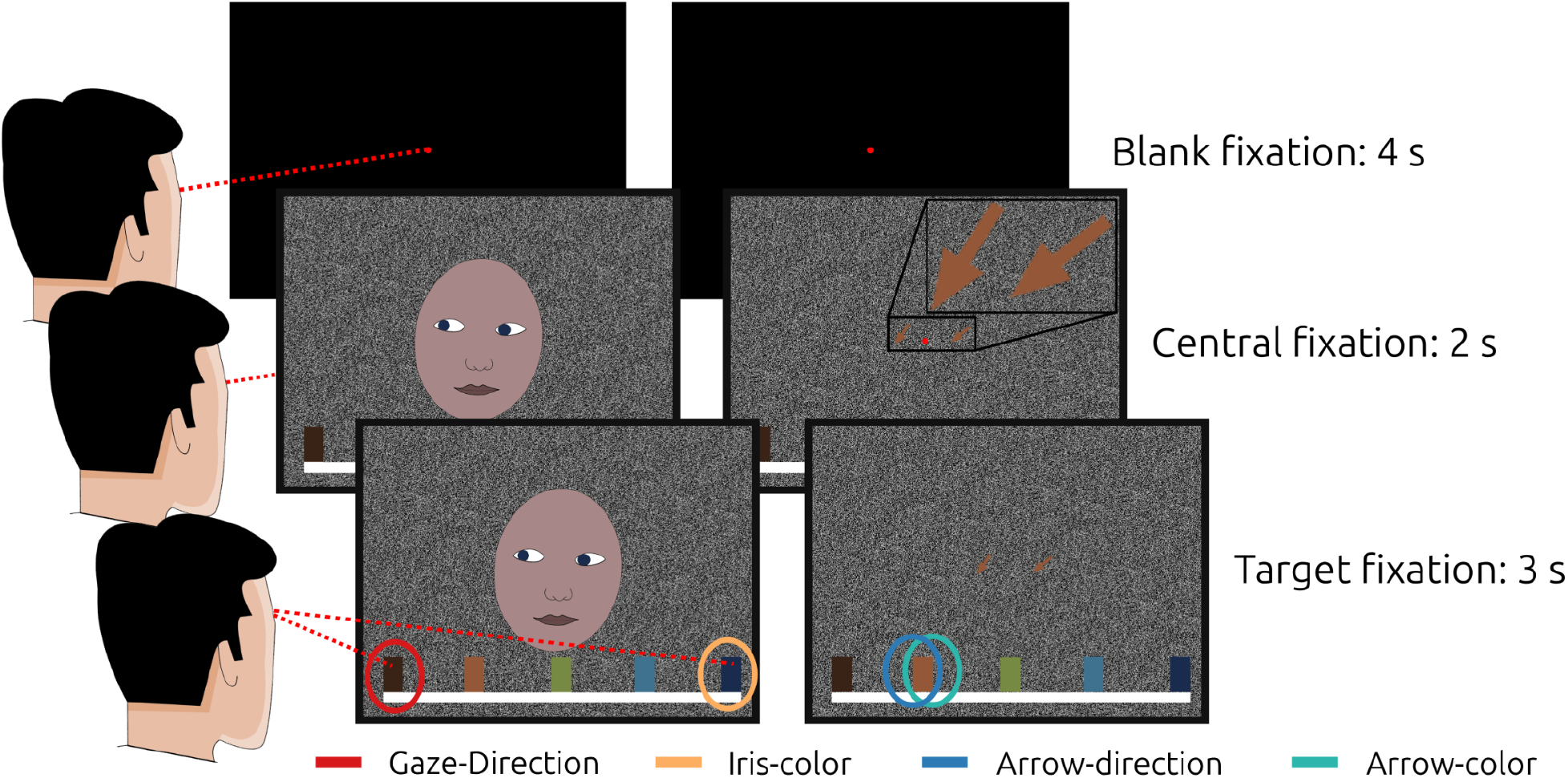
Illustration of the paradigm. The stimulus used in the experiment has been a photograph of a real person, here replaced by a drawing due to privacy reasons. The left and right columns display the sequence of events during a gaze-following/ iris-color mapping and an arrow-direction/ arrow-color mapping trial, respectively. Each trial started with a blank fixation period followed by the presentation of the cue. In each individual trial the demonstrator face and the arrows look/ point towards a randomly selected target and the color of the iris and the arrows, chosen independently of each other, matched one of the targets. After cue presentation the participants had to wait with their response until the central fixation dot disappeared.

## Results

### Contrasts

At a threshold of p < 0.001 (uncorrected) (Figure 2, top) contrasting *gaze-direction following* with *iris-color mapping* (regions encircled in red in Figure 2) did not yield activity overlapping with the GFP as described by Marquardt et al. (2008) (from now on, we refer to the GFP ROI as defined by Marquardt et al. as *m*GFP – the region encircled in pink in Figure 2).^4^ Contrasting *arrow-direction following* with *arrow-color mapping* (regions encircled in blue in Figure 2) yielded activity in the left hemisphere that overlaps with the *m*GFP therein. The coordinates of the local maximum of this patch are (x, y, z) = (−54, -72, 6). The first 5 neurosynth^13^ associations for this location are: *motion, videos, v5, mt, visual*.

**Figure 2.**
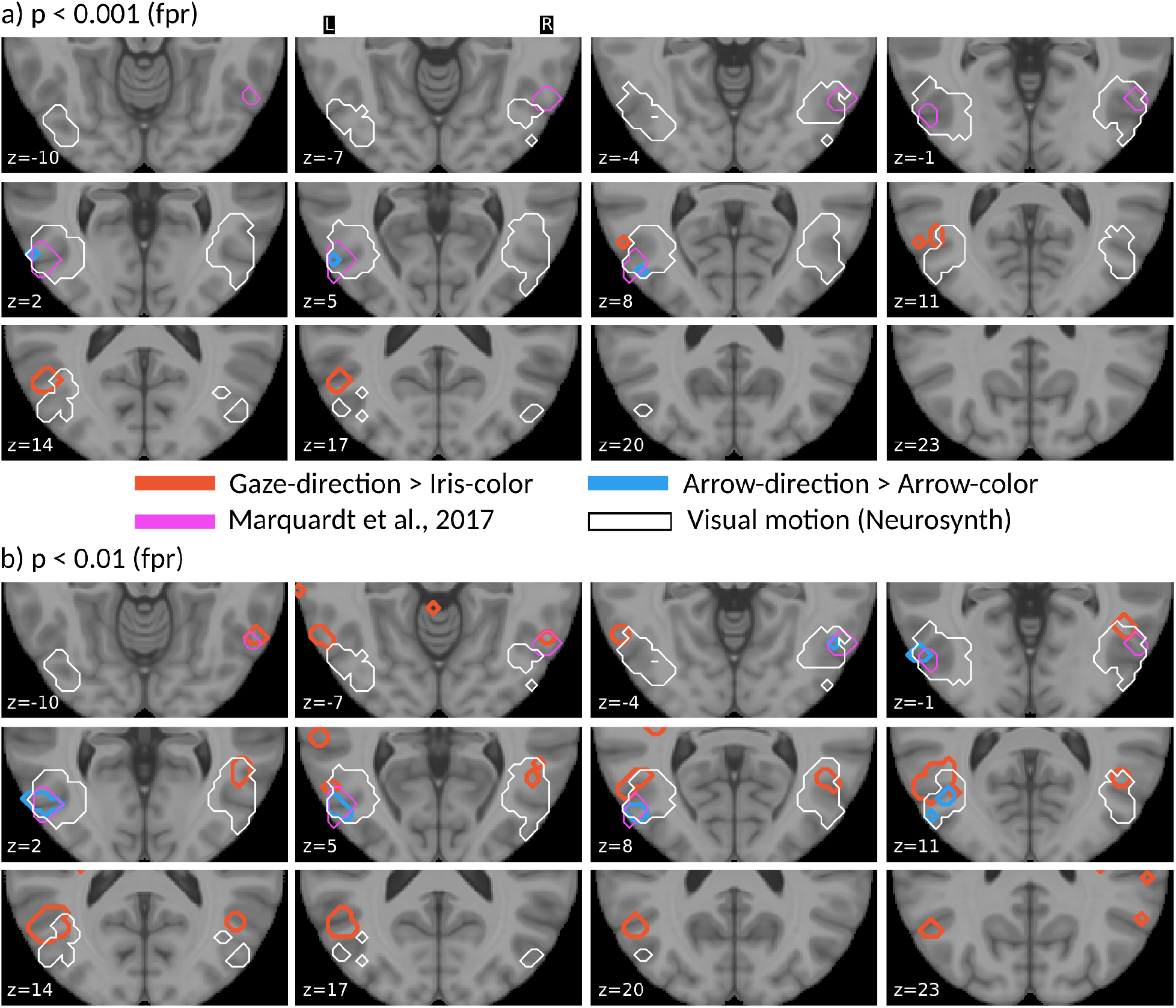
Panel a) and b) both show the same contrasts/ ROIs at two threshold levels. White encircled is the *visual motion* ROI^13^, pink encircled is the *m*GFP^4^. Blue encircled areas belong to the arrows contrast and red encircled areas to the gaze contrast at the respective statistical thresholds.

At this threshold the *gaze-iris* contrast yielded activity dorsal to the *m*GFP beginning to emerge around z = 8 and extending towards z = 18 (encircled red in Figure 2). The local maximum of this patch is located at (x, y, z) = (−51, -63, 15). The first five associations for this location are *action observation, intentions, mentalizing, social, temporoparietal*.

The *m*GFP as well as the *arrows* contrast are falling within the region of the *visual motion* ROI (regions encircled in white in Figure 2.) given by the neurosynth database. The *gaze-iris* contrast is slightly more frontal to this ROI.

Liberalizing the threshold to p < 0.01 (uncorrected) (Figure 2, bottom) additionally gave rise to patches close to or overlapping with the *m*GFP for the *gaze-iris* in both hemispheres (peak coordinates: (x, y, z) = (−60, -60, -3) and (x, y, z) = (57, -60, -6)). Neurosynth associations to these two locations are *word form, judgment task, interactive, semantics, timing* (left) and *visual, unfamiliar, objects, multisensory, visually* (right). Further a patch dorsal to the *m*GFP emerged in the right hemisphere at this threshold for this contrast. This patch had one ventral peak at (48, -54, 0) and a dorsal peak at (48, -58, 12) which closely matched the dorsal activity in the left hemisphere at (−51, -63, 15) already visible at the more conservative threshold of p < 0.001. The neurosynth associations for (48, -54, 0) are *unfamiliar, motion, social interactions, gestures, preparatory* and for (48, -58, 12) *motion, video clips, gaze, action observation*.

At the more liberal threshold the activity patch at (−54, -72, 6) associated with *arrow-direction following* was more extended and matched the *m*GFP, especially in the left hemisphere. In the right hemisphere a small activity patch emerged at this threshold at (51, -63, -3) falling into the area of the *m*GFP, as well. The neurosynth associations for (51, -63, -3) are *visual, motion, objects, movements, action observation*.

Table 1 lists the peak coordinates as well as the neurosynth associations of the activations found in this study as well as the GFP locations reported in two preceding studies on the GFP.

**Table 1.**
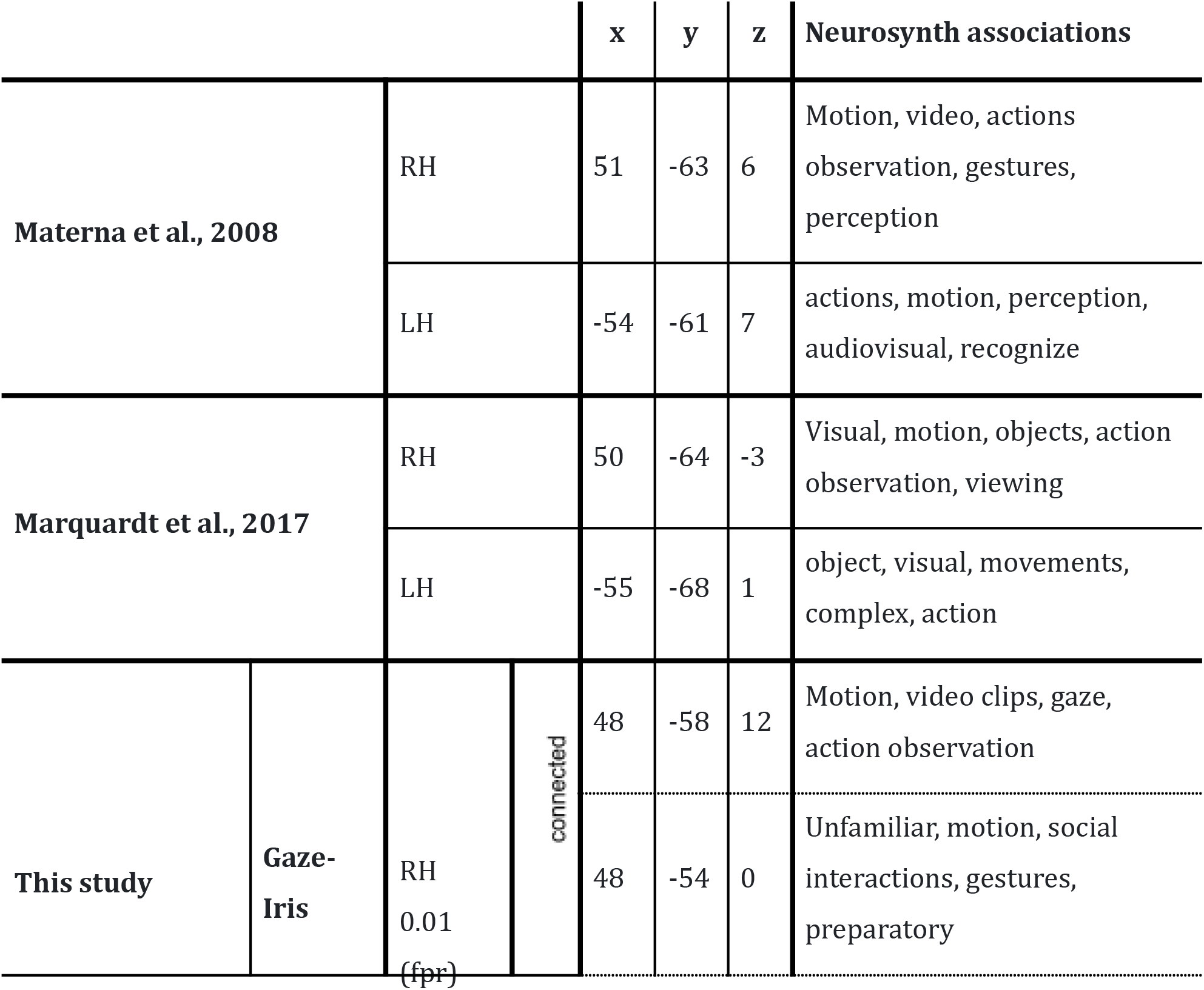
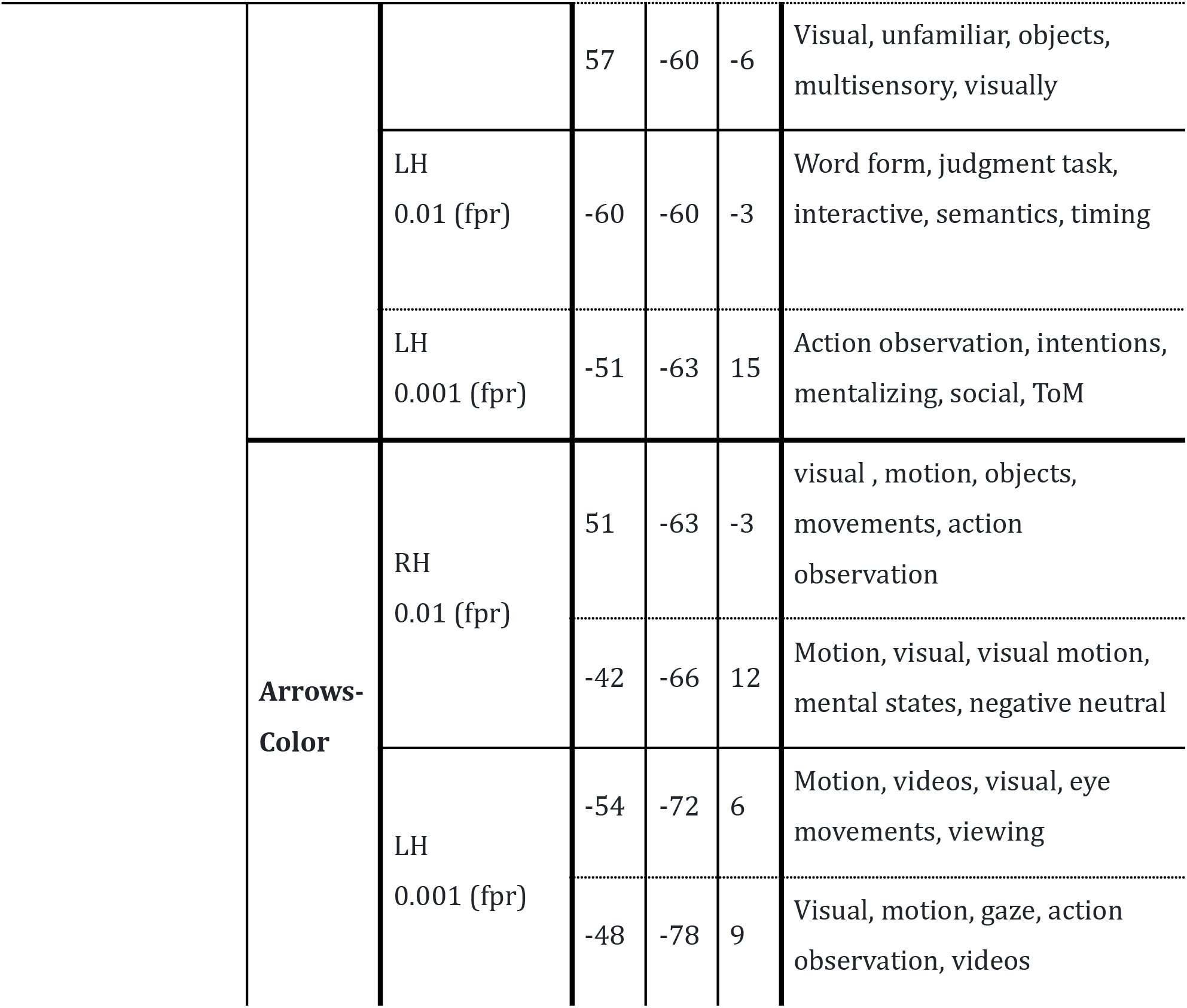
List of locations of the GFP reported in earlier studies as well as locations in its vicinity found in this study. Coordinates are in MNI space. For each location we report the first five functional associations provided by neurosynth.org^13^.

### HRF estimates

Figure 2 shows the estimated HRFs and the 95% CIs for each condition for the *m*GFP ROI and the the ROI based on a search for *visual motion* in the neurosynth database. The HRFs for the *m*GFP show a statistically significant (95% CIs do not include the activation level of 0) deflection around 5 sec after cue onset in both hemispheres in all conditions but the *iris-color* condition. Though the difference is not statistically significant at the 5% level, the activation in the *arrow-direction* condition is nearly twice as large as in the *gaze-direction* following condition in the left hemisphere. This difference is not present in the right hemisphere.

In the *visual motion* ROI the activation level is overall smaller than in the *m*GFP but still significant at the 5% level (see inlets Figure 2, bottom) for all conditions but the iris-color condition in both hemispheres at around 5 sec.

## Discussion

In this study we asked if the GFP – a brain area described in several studies as a domain-specific module dedicated to the processing of others’ gaze direction ^1–5,8^ - is indeed domain-specific or if it is also active when the participants use arrows instead of the gaze of a demonstrator to identify a target object. We used the same portraits as stimuli that were used in previous experiments based upon which the GFP was originally defined but had to introduce a modification to the temporal structure of the task to allow a direct comparison between the gaze and the arrow condition. In the original version of the task that was used in human studies^1,4,14^ as well as in studies with NHP^8,15^ each trial started with a frame displaying the demonstrator looking straight ahead towards the participant, followed by a second frame displaying the demonstrator looking towards the target object. Whenever two consecutive frames feature a coherent shift of pixels that does not shatter the correspondence of the depicted scene in the two frames, the visual system interprets the sequence as motion^9,10^. In the current study the initial frame was replaced by a black screen which was directly followed by the frame displaying the demonstrator looking towards the target object. In light of this modification, the first question here was whether we can replicate the GFP despite the absence of apparent motion in the form of the gaze shift. We have to answer that in the negative. Even though the sample size and the MRI scanner used in this study were the same as in previous studies, at the same statistical threshold used by Marquardt et al. and even at more liberal thresholds we could not detect any activation for the *gaze-direction* minus *iris-color* contrast that coincided with the GFP^4^. However, due to a lack of a positive control for the role of apparent motion we cannot confirm that it is indeed its lack causing the failure to reproduce the GFP. A study specifically designed to test this possibility is in progress^16^.

Even at the statistically most liberal threshold that we applied (*p* < 0.01, *uncorrected*) there was only a small patch related to the contrast *gaze-direction following* minus *iris-color mapping* partially overlapping with the *m*GFP (Figure 2b, Table 1; with “*m*GFP” we denote the GFP ROI as delineated based on the data of Marquardt et al.^4^). Surprisingly, however, the contrast *arrow-direction following* minus *arrow-color* mapping yielded activity patches overlapping with the *m*GFP for both statistical thresholds (see Figure 2 and Table 1). This area of relatively stronger activity during *arrow-direction following* compared to *arrow-color mapping* as well as the *m*GFP fall within parts of a ROI that is associated with the search term *visual motion* in the neurosynth.org^13^ database (see Figure 2, white encircled areas). When searching for the functional associations with the locations activated in the arrow contrast, Neurosynth then also provides *visual, motion, video, eye movements*, etc., as expected (see Table 1).

Activity only detectable for the gaze but not the arrow contrast at *p* < 0.001 (*uncorrected*) can be found dorsal to the *m*GFP in the left hemisphere with its center at x, y, z = -51, -63, 15 (MNI). Anatomically this location corresponds to BA39 or the temporoparietal junction (TPJ). Searching for functional associations of this location at neurosynth.org^13^ yields *action observation, intention, mentalizing, social, ToM* as the first five results (see Table 1). This location was not reported by Marquardt et al.^4^ or Materna et al.^1^ to be activated during gaze following but fits to the idea that perceiving the other’s gaze evokes the assignment of intentions^6,17–19^.

These results provide a surprising picture of the GFP. First, a gaze stimulus lacking apparent motion does not yield a relatively stronger activation when compared to the color-mapping condition at the *m*GFP location. Second, however, arrows pointing towards the target, despite lacking apparent motion as well, yield detectable activity confined to a region very well overlapping with the *m*GFP when contrasted with the color mapping condition. Note, however, that the more conservative threshold of p < 0.001 is not corrected for multiple comparisons, as well.

To capture the activity within the *m*GFP ROI beyond the contrasts and compare it to the activity within the *visual motion* ROI, we modeled the hemodynamic response functions (HRFs) for each of the conditions and hemispheres individually (see Figure 3). We found that all conditions but the *iris-color mapping* condition yielded a significant activation in all ROIs. Moreover, a comparable pattern is apparent in all ROIs, with the *arrow-direction* condition featuring the strongest activation (blue curves in Figure 3). However, there is no statistically significant difference between any condition as the 95% CIs overlap at all time points. Comparison of the general activity patterns obtained for both ROIs shows their similarity, which is not surprising since the *m*GFP overlaps nearly completely with the more lateral part of the visual motion ROI. The overall smaller amplitude of the HRF for the latter ROI is likely due to the fact that the *visual motion* ROI was about five times larger than the *m*GFP, and therefore most probably comprised voxels which added task-independent signals.

**Figure 3.**
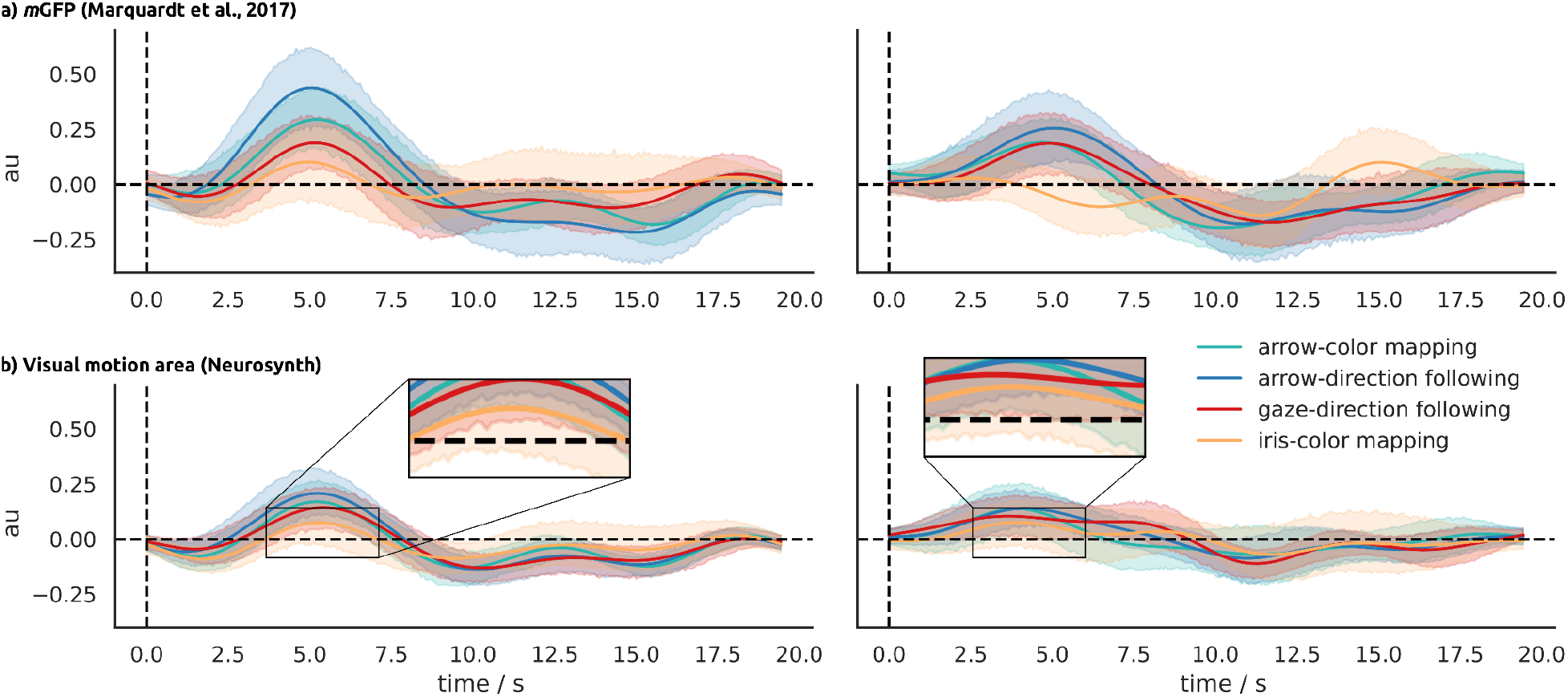
In (a) and (b), the HRF estimates for the four different conditions in the left- and right-hemispheric *m*GFP and *visual motion* ROI, respectively, are shown. Shaded areas represent the 95% CIs. Especially in the left *m*GFP the dominance of activity related to the arrow-direction following condition is obvious.

But what is it then that triggers the activation of the mGFP, given the absence of apparent motion? Taking the results of the contrast analysis and HRF estimates together, we can think of three possible explanations. First, it might be a nonspecific response within the visual system to the switch from a blank screen to the stimulus image. Indeed, it was shown that the BOLD activity in the early visual cortex is positively correlated with scene complexity^20^. Unfortunately, it appears unclear whether the apparently higher visual complexity/ size of the face stimulus compared to the arrows should lead to a stronger activation in the visual system. Or whether this is to be expected from the less conspicuous arrows, which are for this very reason - corresponding to a de facto higher complexity - more difficult to recognize, thus resulting in a higher workload. On that ground we cannot exclude this possibility. The second possibility is that the location corresponding to the mGFP ROI is relevant for the processing of the behaviorally relevant orientation of objects - independent of the type of object. However, we are not aware of any studies that would directly support this hypothesis. Some studies suggest neural tuning to object orientations in area V4^21^ as well as in parietal and frontal areas^22^. But because their paradigms and ours are very different, it is not clear how the results can be related to each other. As a third possibility, we propose that the observed activity patterns can be attributed to an effect described as *implied motion*. Both psychophysical^23^ as well as fMRI studies^24,25^ have demonstrated that static images of moving objects elicit experimental effects that are known from the perception of visual motion (motion adaptation). How could this effect explain our results, given the fact that the stimulus images used here do not depict moving objects? Guterstam and colleagues published a series of studies^26–28^ (compare our Comment Letter^29^) in which they demonstrated that viewing a static image of a cartoon face looking towards an object is indeed able to evoke motion-adaptation in the viewer. In an fMRI study^30^ they furthermore showed that these behavioral effects are accompanied by activity within the human MT+ complex. However, in their data they find that these effects are unique to gaze and are not present if the stimulus is an arrow, contradicting our results here. A study by Lorteije et al.^31^, however, presented evidence that the effects typically attributed to implied motion can be attributed to low-level features of the stimuli like orientation and size. Thus, the effect we find here for arrows seems plausible, though the interpretation would change accordingly. In fact, the results from Lorteije et al. are related to the second explanation (orientation tuning) that we describe above.

Other authors have used arrows as attentional cues and compared them to gaze in spatial cueing tasks inspired by Posner^32^ before us. However, due to the differences in the tasks, no conclusive comparisons can be made with our results, yet they help to contextualize our findings. For example Hietanen et al.^33^ found that gaze and arrows, when used as central endogenous cues, elicit the same behavioral effects, but the BOLD activations differ. Activity related to arrows was much more widespread involving all cortical lobes while activity related to gaze was confined to the inferior and middle occipital gyri, however, in some distance to the *m*GFP. A study by Callejas and colleagues^34^, on the other hand, reported that largely the same neural networks encode gaze and arrow cues but that some differential modulations occurred in parts of the intraparietal sulcus and in the MT+ region. Interestingly they found that the MT+ region exhibited effects only in relation to arrow cues. In that both studies report a stronger BOLD activation associated with arrow cues compared to gaze cues, our results are consistent. However, other than in Callejas’ study, we also find a significant, albeit smaller, activation in the MT+ region in response to static gaze cues. As an explanation for the greater activity associated with the arrow stimuli, the authors suggest the greater automaticity of responses to gaze stimuli, as well as that the processing of arrow stimuli could be more demanding. Given that the gaze of others is ubiquitous from birth and plays a crucial role in ontogeny the processing of faces and gaze is likely to be highly optimized resulting in smaller signal changes when investigated using fMRI. There are two neuropsychological cases that are of great interest, as well. The one is reported by Akiyama et al.^35^ and consists of an impairment of using gaze but not arrows as a spatial cue after a lesion to the (entire) right superior temporal gyrus. This case confirms that the superior temporal region, which was reported to be involved in face perception^36–38^, especially in the perception of changeable aspects of faces such as eye and mouth movements^39,40^, is an integral component and that its lack leads to impairments in tasks in which eye recognition plays a crucial role. The other case is reported in two studies by Vecera and Rizzo^41,42^ and points towards another brain region that may be crucial for the perception and interpretation of gaze cues, as well - the frontal lobe. They report the case of a patient with frontal lobe damage who had lost the ability to direct attention to peripheral locations volitionally through endogenous cues such as words and gaze, but could still attend automatically to exogenous cues. Based on this they suggest an association hypothesis that the gaze of another person is understood, analogous to words, through associations of gaze directions with locations in space.

All this leaves a situation involving a number of perspectives impossible to reconcile at this point. In sum, the present study provides evidence that the activity within GFP, as reported in previous studies, may largely depend on the presence of (apparent and/ or implied) motion in the stimuli since its reproduction failed in the absence of visual motion. But to which degree low level visual motion is sufficient to trigger the GFP cannot be answered here.

## Methods

### Paradigm & Setup

The paradigm consisted of 2 stimuli types with 2 conditions, each. One stimulus type consisted of a photo of a face and the other of 2 arrows that were displayed on a computer monitor. Below the stimuli, 5 differently colored squares were shown, which served as gaze targets. Each trial started with a black screen with a red fixation dot in the middle which was shown for 4 sec. After this either the face or the arrows were displayed, still with the red fixation dot in the center of the monitor. The face looked in the direction of one of the squares, the iris-color matched one of the colors of the squares, the arrows pointed towards one of the squares and their color matched one of the squares, as well. After 2 more seconds the fixation dot disappeared which was the go-cue telling the participants that they now have to shift their own visual focus onto the target square defined by the current experimental condition. Conditions were organized in blocks such that the participants always had to identify the target square by the same stimulus features for 20 consecutive trials. Which stimulus feature mattered was announced by a written instruction before each block. Four blocks made up a run within which each condition occured in randomized order. In total 4 runs had to be completed, such that each condition was represented by 80 trials.

During the experiment participants lay in the scanner and viewed the stimulus monitor via a mirror system. The distance between the participants eyes and the monitor was ∼190 cm and it covered ∼20° of the field of view in the horizontal and ∼12° in the vertical. Additionally, eye tracking data was recorded during the fMRI experiment. In a separate session participants were familiarized with the task outside of the scanner.

### Data collection

MR images were acquired in a 3T scanner (Siemens Magnetom Prisma) with a 20-channel phased array head coil. The head of the subjects was fixed inside the head coil by using plastic foam cushions to avoid head movements. An AutoAlign sequence was used to standardize the alignment of images across sessions and subjects. A high-resolution T1-weighted anatomic scan (MP-RAGE, 176 × 256 × 256 voxel, voxel size 1 × 1×1 mm) and local field maps were acquired. Functional scans were conducted using a T2^*^-weighted echo-planar multibanded 2D sequence (multiband factor = 2, TE = 35ms, TR=1500ms, flip angle = 70°) which covers the whole brain (44 × 64 × 64 voxel, voxel size 3 × 3 × 3 mm, interleaved slice acquisition, no gap).

### Preprocessing

Results included in this manuscript were preprocessed using fMRIPrep 1.5.2^43,44^.

#### Copyright Waiver

The below boilerplate text was automatically generated by fMRIPrep with the express intention that users should copy and paste this text into their manuscripts unchanged. It is released under the CC0 license.

#### Anatomical data preprocessing

The T1-weighted (T1w) image was corrected for intensity non-uniformity (INU) with N4BiasFieldCorrection^45^, distributed with ANTs 2.2.0^46^, and used as T1w-reference throughout the workflow. The T1w-reference was then skull-stripped with a Nipype implementation of the antsBrainExtraction.sh workflow (from ANTs), using OASIS30ANTs as target template. Brain tissue segmentation of cerebrospinal fluid (CSF), white-matter (WM) and gray-matter (GM) was performed on the brain-extracted T1w using fast^47^ (FSL 5.0.9). Brain surfaces were reconstructed using recon-all^48^ (FreeSurfer 6.0.1), and the brain mask estimated previously was refined with a custom variation of the method to reconcile ANTs-derived and FreeSurfer-derived segmentations of the cortical gray-matter of Mindboggle^49^. Volume-based spatial normalization to one standard space (MNI152NLin2009cAsym) was performed through nonlinear registration with antsRegistration (ANTs 2.2.0), using brain-extracted versions of both T1w reference and the T1w template. The following template was selected for spatial normalization: ICBM 152 Nonlinear Asymmetrical template version 2009c^50^ (TemplateFlow ID: MNI152NLin2009cAsym).

#### Functional data preprocessing

For each of the 4 BOLD runs found per subject (across all tasks and sessions), the following preprocessing was performed. First, a reference volume and its skull-stripped version were generated using a custom methodology of fMRIPrep. A deformation field to correct for susceptibility distortions was estimated based on a field map that was co-registered to the BOLD reference, using a custom workflow of fMRIPrep derived from D. Greve’s epidewarp.fsl script and further improvements of HCP Pipelines^51^. Based on the estimated susceptibility distortion, an unwarped BOLD reference was calculated for a more accurate co-registration with the anatomical reference. The BOLD reference was then co-registered to the T1w reference using bbregister (FreeSurfer) which implements boundary-based registration^52^. Co-registration was configured with six degrees of freedom. Head-motion parameters with respect to the BOLD reference (transformation matrices, and six corresponding rotation and translation parameters) are estimated before any spatiotemporal filtering using mcflirt^53^ (FSL 5.0.9). BOLD runs were slice-time corrected using 3dTshift from AFNI 20160207^54^. The BOLD time-series were resampled to surfaces on the following spaces: fsaverage5. The BOLD time-series (including slice-timing correction when applied) were resampled onto their original, native space by applying a single, composite transform to correct for head-motion and susceptibility distortions. These resampled BOLD time-series will be referred to as preprocessed BOLD in original space, or just preprocessed BOLD. The BOLD time-series were resampled into standard space, generating a preprocessed BOLD run in [‘MNI152NLin2009cAsym’] space. First, a reference volume and its skull-stripped version were generated using a custom methodology of fMRIPrep. Several confounding time-series were calculated based on the preprocessed BOLD: framewise displacement (FD), DVARS and three region-wise global signals. FD and DVARS are calculated for each functional run, both using their implementations in Nipype (following the definitions by Power et al. 2014^55^). The three global signals are extracted within the CSF, the WM, and the whole-brain masks. Additionally, a set of physiological regressors were extracted to allow for component-based noise correction (CompCor^56^). Principal components are estimated after high-pass filtering the preprocessed BOLD time-series (using a discrete cosine filter with 128s cut-off) for the two CompCor variants: temporal (tCompCor) and anatomical (aCompCor). tCompCor components are then calculated from the top 5% variable voxels within a mask covering the subcortical regions. This subcortical mask is obtained by heavily eroding the brain mask, which ensures it does not include cortical GM regions. For aCompCor, components are calculated within the intersection of the aforementioned mask and the union of CSF and WM masks calculated in T1w space, after their projection to the native space of each functional run (using the inverse BOLD-to-T1w transformation). Components are also calculated separately within the WM and CSF masks. For each CompCor decomposition, the k components with the largest singular values are retained, such that the retained components’ time series are sufficient to explain 50 percent of variance across the nuisance mask (CSF, WM, combined, or temporal). The remaining components are dropped from consideration. The head-motion estimates calculated in the correction step were also placed within the corresponding confounds file. The confound time series derived from head motion estimates and global signals were expanded with the inclusion of temporal derivatives and quadratic terms for each^57^. Frames that exceeded a threshold of 0.5 mm FD or 1.5 standardized DVARS were annotated as motion outliers. All resamplings can be performed with a single interpolation step by composing all the pertinent transformations (i.e. head-motion transform matrices, susceptibility distortion correction when available, and co-registrations to anatomical and output spaces). Gridded (volumetric) resamplings were performed using antsApplyTransforms (ANTs), configured with Lanczos interpolation to minimize the smoothing effects of other kernels^58^. Non-gridded (surface) resamplings were performed using mri_vol2surf (FreeSurfer).

Many internal operations of fMRIPrep use Nilearn 0.5.2^59^, mostly within the functional processing workflow. For more details of the pipeline, see the section corresponding to workflows in fMRIPrep’s documentation.

### Analysis

#### Contrasts

For each participant we computed a GLM (general linear model) for the combination of all runs (first-level) to obtain the respective β-images for each condition using the Python package nilearn^60^. For modeling, we aligned the *onsets* of each trial to the onsets of the spatial/ color cue (4 sec after the actual trial onset, starting with the blank screen) and set the stimulus duration to 0, i.e. we modeled the relevant stimulus component as a single event since it is not changing over time. To mitigate the effects of motion artifacts and other noise sources, the nuisance regressors global_signal, csf, white_matter, trans_x, trans_y, trans_z, rot_x, rot_y, rot_z and their respective first derivatives estimated by fMRIPrep were included in the design matrices. As the model for the HRF we used the *glover + derivative + dispersion* provided by nilearn and included a polynomial drift model of order 3 to remove slow drifts in the data. Further, we masked the data with the average across run’s mask images provided by fMRIPrep and applied smoothing with *fwhm* = 8 mm.

For each participant the resulting first-level *β*-images will be used to compute the contrasts *gaze-direction – iris-color* and *arrow-direction – arrow-color*. The resulting *effect-size* images (nilearn terminology) of each contrast were fed into second-level analyses that were fitted as one-sample t-tests. The resulting second-level contrasts were thresholded with p<0.001 (fpr) and p<0.01 (fpr). All coordinates reported in this manuscript refer to the MNI space. For each of the reported coordinates a neurosynth search was conducted and we report the first 5 associations.

#### ROI definition & HRF estimation

To define the GFP ROI we extracted the activity patch of both hemispheres as reported by Marquardt et al^4^ and converted them into mask images. To define a ROI corresponding to brain areas that process visual motion we searched the neurosynth database (*Term-based meta-analysis*) using the term *visual motion*. After downloading the result (*association test*) we extracted the largest components in both hemispheres spanning the posterior temporal cortex and converted them into mask images. The left GFP consists of 76 voxels, the right of 61 voxels and the visual motion ROIs consist of 377 (left) and 306 (voxels). All ROIs are plotted along with the contrasts described above in Figure 2.

For each participant and ROI, we estimated the hemodynamic response of each condition. To do so, the BOLD signals of each ROI (averaged across voxels) were extracted. Prior to signal extraction the BOLD images were denoised using the same nuisance regressors as used to fit the GLMs described above. To recover the condition specific hemodynamic responses we deconvolved the signals using the nideconv package^61^. We applied the *fourier* basis set with 9 regressors over a period of 19.5 sec starting from spatial/ color cue onset, again omitting the blank screen period in the beginning of each trial. Individual models were fitted using the standard settings of nideconv’s fitting method. The first-level response estimates were then fed into the group-level model using nideconv’s GroupResponseFitter functionality using the same settings as for the first-level. As a result we obtain the estimated HRFs together with the 95% credible intervals (CI) of the estimates. Periods in which the CIs do not include an activation level of 0 (au) are considered to be statistically different from 0 at the 5% level.

